# Alternative splicing of *PIF4* regulates plant development under heat stress

**DOI:** 10.64898/2025.12.17.694898

**Authors:** María Niño-González, Benjamin Alary, Dóra Szakonyi, Tom Laloum, Paula Duque, Guiomar Martín

## Abstract

The Phytochrome-Interacting Factor 4 (PIF4) is a key player in the integration of multiple internal and external stimuli to optimize different aspects of plant development. While both the DNA encoding this transcription factor and its protein are known to be under tight control, no regulation at the RNA level has been previously reported. Our genomic analysis revealed that the exon/intron structure of the basic Helix-Loop-Helix (bHLH) DNA binding domain of PIF4 is conserved and pointed to skipping of an exon in this region specifically in response to heat stress. We then showed that this alternative splicing event downregulates PIF4 function under heat, which in etiolated seedlings induces photomorphogenic-related traits. Our results disclose a role for PIFs in plant responses to heat and reveal a new regulatory layer for the control of PIF4 function, underscoring the critical role of posttranscriptional regulatory processes in the molecular integration of environmental cues.

## Introduction

Phytochrome Interacting Factors (PIFs) belong to the basic Helix-Loop-Helix (bHLH) family of transcription factors. The bHLH protein domain consists of two segments: the basic region, required for DNA binding, and the helix-loop-helix, responsible for hetero and homodimerization (Toledo-Ortiz et al., 2003). PIFs are also characterized by the presence of a protein domain that interacts with photoactivated phytochromes (Favero, 2020). This interaction promotes the degradation of PIFs in the light and is crucial for their role as major regulators of light-regulated biological processes (Leivar and Monte, 2014). In etiolated seedlings, which germinate and develop in subterranean darkness, PIFs are active and repress photomorphogenic features, such as chlorophyll biosynthesis, cotyledon expansion, and repression of hypocotyl elongation (Leivar et al., 2008; Shin et al., 2009).

Besides their well-known function in adjusting seedling development to light, PIFs, particularly PIF4, have over the past years also been implicated in the regulation of different biological processes such as immunity (Gangappa et al., 2017), morphological adaptations to high ambient temperatures (Koini et al., 2009), stomatal development (Casson et al., 2009), leaf senescence (Sakuraba et al., 2014), freezing tolerance (Lee and Thomashow, 2012), salt tolerance (Wang et al., 2025), anthocyanin biosynthesis (Liu et al., 2015), or fatty acid biosynthesis (Liao et al., 2025). PIF4 has thus emerged as a key integrator of multiple external and internal signals to optimize plant development (Choi and Oh, 2016; Lucyshyn and Wigge, 2009).

Several studies have described the molecular mechanisms that control *PIF4* gene expression, protein levels, and activity (Leivar and Quail, 2011; Paik et al., 2017; Pham et al., 2018), but its posttranscriptional regulation has never been characterized. Here we show that alternative splicing, a posttranscriptional process generating multiple mRNAs from the same gene, produces two different *PIF4* transcripts specifically in response to heat stress.

Temperature deviations from the optimal range significantly impact plant development and survival. Increases in temperature are classified as either high ambient temperature or excessively hot temperatures. High ambient temperature is typically 5-6ºC above the optimum temperature (22ºC for *Arabidopsis thaliana*), while excessively hot temperatures exceed this range (Li et al., 2018). These distinct temperature ranges activate independent signaling pathways, leading to different physiological outcomes. Warm temperatures induce thermomorphogenesis, which generally promotes growth and development in a PIF4-dependent manner (Quint et al., 2016). Conversely, excessively hot temperatures trigger stress-responsive pathways aimed at adjusting growth and physiology to mitigate the negative effects of heat (Kan et al., 2023). To date, the role of PIFs in temperature signaling has centered on thermomorphogenesis, with only a few studies having explored the role of PIF proteins in heat stress responses (Li et al., 2021; Yang et al., 2022). Intriguingly, our results reveal that heat stress induces photomorphogenic features in etiolated seedlings and that this developmental response is mediated by an alternative splice form of *PIF4*.

## Results

### *PIF4* is alternatively spliced in response to heat stress

Our analysis of the exon and intron positioning in all 15 members of the XV subfamily of *Arabidopsis thaliana* bHLH transcription factors that includes PIFs (Toledo-Ortiz et al., 2003) showed that, with the sole exception of the less conserved member HFR1, the intron/exon distribution of the bHLH domain is maintained. Its 150-bp long sequence is distributed among three exons, following an invariable proportion: 22%, 44% and 34% (Figure 1A). In addition, the middle exon containing the largest section of the bHLH domain is always 66-bp long and surrounded by phase 0 introns, which preserve codon identities and therefore the reading frame (Figure 1A). Hence, alternative splicing of this exon will produce protein isoforms differing only in the bHLH region (Supplemental Figure 1). Given the genomic particularities of the bHLH middle exon in these genes, we investigated its splicing regulation by quantifying its PSI (Percent Spliced In; percentage of transcripts that include the exon) in publicly available *Arabidopsis thaliana* RNA-seq samples covering several environmental conditions and tissues at different developmental stages (Supplemental Table 1). This analysis revealed skipping of the exon exclusively in two genes: *PIF4* and *PIF6* (Figure 1B and Supplemental Table 1). In the case of *PIF6*, although exon skipping occurs in nearly all tissues and conditions tested, this alternative splicing event has only been shown to be functional in seeds and embryos (Penfield et al., 2010), where *PIF6* is highly expressed (Supplemental Figure 2). Strikingly, we found that the *PIF4* gene undergoes alternative splicing in this particular exon, exon 5, exclusively when plants are under heat stress (Figure 1B and Supplemental Figure 3). Given that the protein arising from this exon skipping event will lack a portion of the bHLH domain (Supplemental Figure 1), we expect heat-induced alternative splicing to reduce the amounts of functional PIF4.

**Figure 1.**
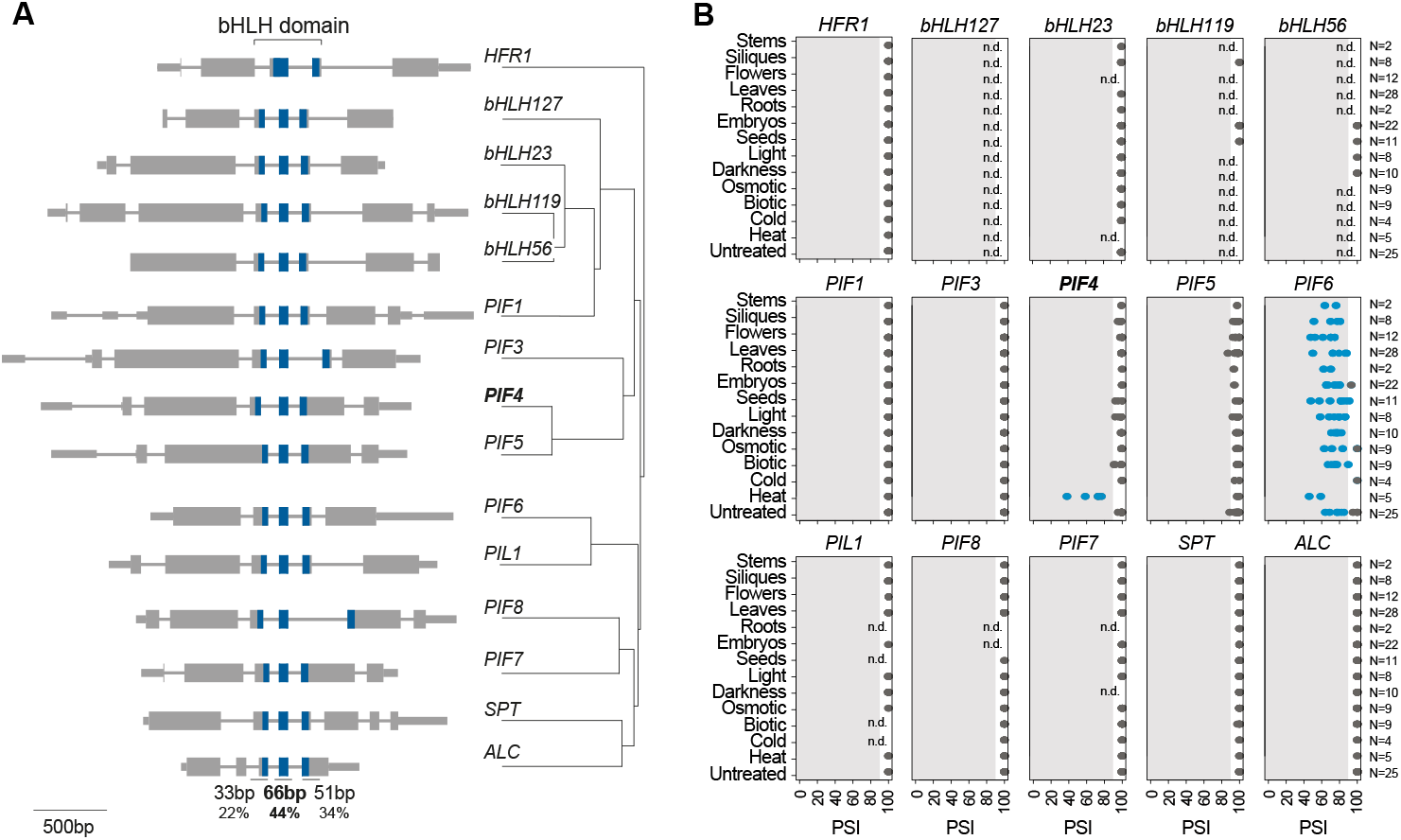
Alternative splicing regulation of the bHLH central exon in the XV subfamily of Arabidopsis *thaliana* bHLH transcription factors. **(A)** Exon and intron gene distribution (left) and phylogeny (right; adapted from Leivar and Quail, 2011) of all members of the XV subfamily of Arabidopsis bHLH transcription factors. The bHLH subdomains are shown in blue. For all genes with the exception of *HFR1*, percentages indicate the proportion of the bHLH domain encoded in each exon. **(B)** PSI (Percent Spliced In) of the bHLH central exon using publicly available RNA-seq data covering several *Arabidopsis thaliana* tissues and environmental conditions (Supplemental Table 1). Blue dots indicate PSI<90, the generally considered cutoff for exon skipping. n.d., not detected.

### Heat stress induces photomorphogenesis in the dark

To study the impact of the exon skipping event in the bHLH domain of *PIF4*, we applied heat stress (37 ºC) to 3-day-old etiolated seedlings, a developmental stage at which PIFs are known to be functional in repressing photomorphogenesis (Leivar et al., 2008; Shin et al., 2009). First, we confirmed that the heat-induced *PIF4* alternative splicing event occurs in etiolated seedlings as well (Figure 2A), also showing that it does not occur in response to light and is sustained along time (Figure 2A and 2B). Remarkably, heat treatment of dark-grown seedlings partially induces photomorphogenesis — cotyledons open and hypocotyl elongation is repressed (Figure 2C and 2D). These morphological changes, although less pronounced, are characteristic of seedlings lacking PIF activity, as is the case with etiolated seedlings transferred to light or dark-grown quadruple *pif1pif3pif4pif5* (*pifq*) mutants (Figure 2B and Supplemental Figure 4) (Leivar et al., 2009, Leivar et al., 2008; Shin et al., 2009). Moreover, we quantified protochlorophyllide (Pchlide), the phototoxic chlorophyll precursor that, when overaccumulated in etiolated seedings, leads to photobleaching upon light exposure. This analysis revealed higher levels of Pchlide and increased photobleaching in wild-type (WT) etiolated seedlings exposed to heat for 24 hours prior to light exposure, a trend also partially phenocopying *pifq* mutants (Figure 2E, 2F and Supplemental Figure 5) (Leivar et al., 2009). These phenotypes are consistent with heat-induced alternative splicing reducing the amounts of functional PIF4. Analysis of *PIF1, PIF3, PIF4* and *PIF5* expression levels under 37ºC demonstrated that none of these genes were transcriptionally downregulated by heat stress (Figure 2G and Supplemental Figure 6), discarding a strong reduction in *PIF* transcript levels as the cause of the observed phenotypes. Heat-stressed etiolated seedlings are phenotypically more similar to higher order *pif* mutants than to *pif4* single mutants (Supplemental Figure 7 and 8) (Leivar et al., 2008; Shin et al., 2009), suggesting that the *PIF4-S* splice form generated by exon skipping may act as a dominant negative rather than being merely inactive. Previous studies have reported that different PIF4 protein isoforms can exert dominant negative effects, inhibiting the activity of other PIFs, and that alternative splicing can produce dominant negative transcription factor isoforms (Gangappa et al., 2017; Kim et al., 2020; Nicolas et al., 2015; Seo et al., 2011). However, direct evidence is needed to confirm that the *PIF4* short isoform functions as a dominant negative, reducing the activity of the long isoform and other PIFs. Nevertheless, because heat-induced phenotypes are milder than those observed in *pifq* seedlings (Figure 2C, 2E and 2F), a substantial fraction of PIFs likely remains functional at 37 ºC. This is consistent with heat-induced alternative splicing affecting only around 50% of transcribed *PIF4* mRNAs (Figure 2A and 2B).

**Figure 2.**
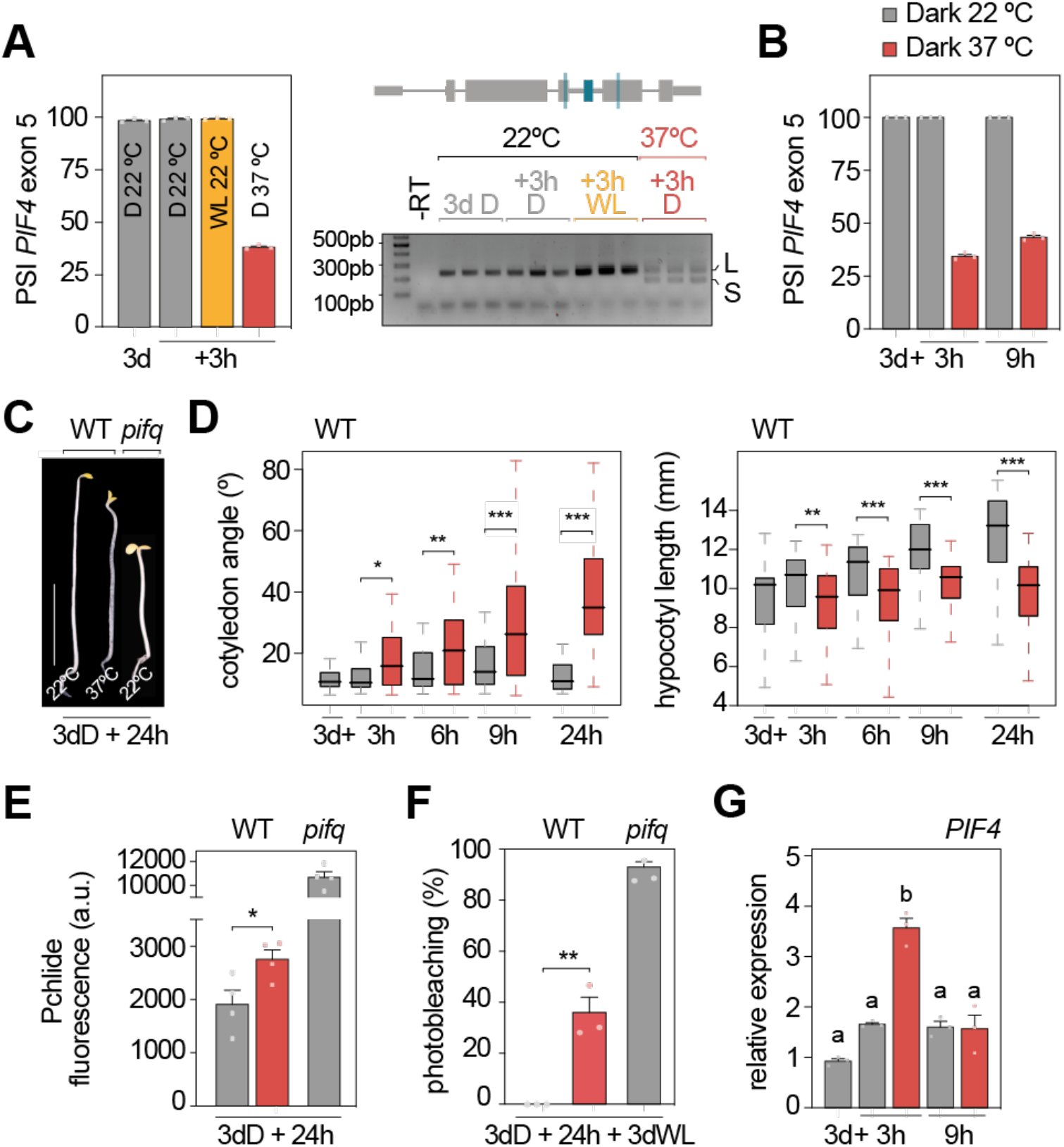
Impact of heat treatment in etiolated seedlings. **(A-B)** PSI (Percent Spliced In) quantification (left) from RT-PCR (right) of the *PIF4* alternatively-spliced exon in seedlings grown in continuous dark for 3 days (d) and then transferred to 37 ºC (red) or white light (WL; yellow), or kept at 22 ºC for 3 (A) or 9 (B) additional hours (h) in darkness (D; gray). Data represent means ± SEM of biological triplicates. -RT, no reverse transcriptase. **(C)** Representative image of 3-day-old wild-type (WT) and *pifq* seedlings subjected or not to heat stress. Scale bar, 5 mm. **(D)** Boxplot representations of the cotyledon aperture (left) and hypocotyl length (right) of at least 35 WT seedlings grown in darkness for 3 days and then transferred to 37 ºC in darkness (red) or kept in the dark at 22 ºC (gray) for the indicated time in hours. Asterisks indicate statistically differences between medians (Mann Whitney test). Fluorescence of protochlorophyllide (Pchlide; 635 nm) **(E**; *n*=4) and percentage of photobleaching **(F**; *n*=3) in heat-stressed (red) or unstressed (gray) etiolated WT and *pifq* seedlings. Asterisks indicate statistically differences between averages (*t*-test). **(G)** RT-qPCR analysis of *PIF4* transcript levels in WT seedlings grown as in (*B*). Values were normalized to *PP2A* and expression levels are expressed relative to the initial time point set at one. Data represent means ± SEM of biological triplicates, and different letters denote statistically significant differences under each condition (Tukey test; *P*<0.05). n.d., not detected. (A, D, E, F) \**P*<0.05, \*\**P*<0.01 and \*\*\**P*<0.001.

### Heat-induced photomorphogenesis depends on *PIF4* alternative splicing

To confirm the implication of *PIF4* alternative splicing in the physiological changes undergone by heat-stressed etiolated seedlings, we quantified the morphological and chlorophyll-related phenotypes of transgenic plants expressing predominantly the long *PIF4* splice form in the *pif4-101* mutant background, (*PIF4p::PIF4-L*; Figure 3A and Supplemental Figure 9). Importantly, both the heat-induced Pchlide accumulation and cotyledon opening were strongly reduced in these lines, while the repression of hypocotyl elongation was maintained (Figure 3B, 3C and Supplemental Figure 10). In agreement with the reduction of Pchlide in *PIF4-L* plants, a non-significant but correlating trend in their photobleaching phenotypes was also observed (Figure 3B). Importantly, *PIF4-L*.*1* expresses the long isoform at levels similar to those of WT plants (Supplemental Figure 9A), ruling out the possibility that the suppression of heat-induced phenotypes (cotyledon opening and Pchlide accumulation) is due to elevated *PIF4* expression levels. In addition, consistent with the comparable alternative splicing levels observed in heat-stressed WT and *pif4* seedlings (Figure 3A and Supplemental Figure 9B), the skipped exon is located upstream of the *pif4-101* mutation (Supplemental Figure 9C), and the phenotypes are also comparable (Figure 3). Similar results were obtained with the other commonly used *pif4* mutant (*pif4-2*; Leivar et al., 2008), which harbors a similar insertion site (Supplemental Figure 9D). Overall, our results substantiate a role for *PIF4* alternative splicing in controlling heat-induced developmental responses in etiolated seedlings. Next, we quantified the transcriptional response to heat stress in etiolated seedlings with different levels of the long *PIF4* splice form (Figure 3A). Interestingly, we found a strong enrichment of heat-regulated genes among those reported as PIFq-regulated or PIFq-bound (Pfeiffer et al., 2014) (Figure 3D). Furthermore, heat-induced transcriptional changes in *pif4* mutants, the genetic background of *PIF4-L* seedlings, were significantly attenuated in these transgenic lines, yet the response remained far from abolished (Figure 3E). This result could be explained by some heat-induced transcriptional changes being fully PIF-independent, as shown in Figure 3D, and others being PIF-dependent but unaffected due to the considerable fraction of PIFs still functional under heat stress. Either scenario would also explain the partial reversion of the heat-induced phenotypic responses observed in *PIF4-L* lines.

**Figure 3.**
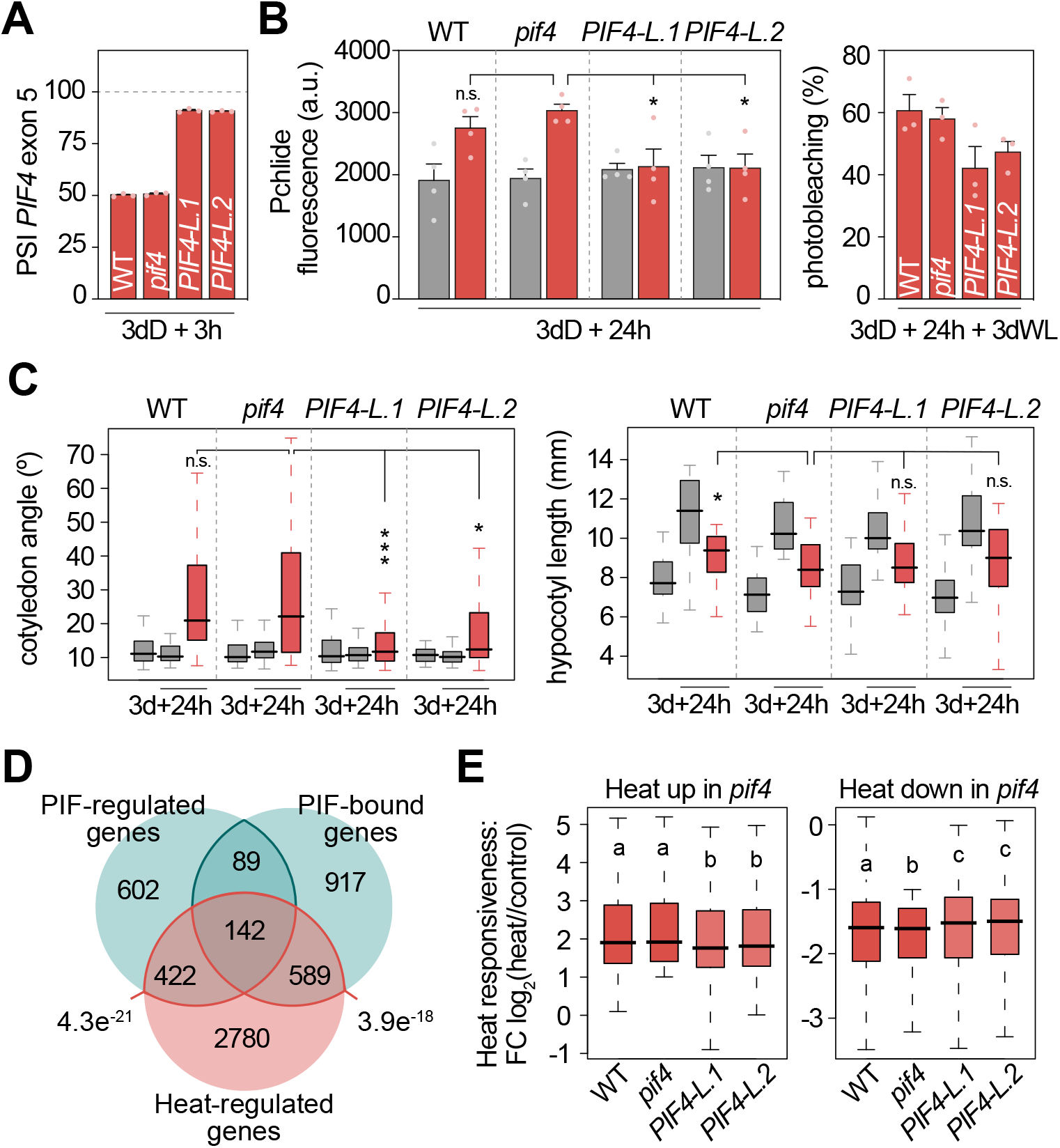
Enhancing expression of the *PIF4* long isoform at 37ºC reduces the impact of heat stress in etiolated seedlings. **(A)** PSI (Percent Spliced In) quantification of the *PIF4* alternatively-spliced exon in wild-type (WT), *pif4* and *PIF4-L* seedlings grown in continuous dark for 3 days (d) and then transferred to 37 ºC for 3 additional hours (h) in darkness. Data represent means ± SEM of biological triplicates. **(B)** Protochlorophyllide (Pchlide; 635 nm) fluorescence (left; *n*=4) and percentage of photobleaching (right; *n*=3) in 3-day-old etiolated WT, *pif4* and *PIF4-L* seedlings transferred to 37 ºC (red) or kept at 22 ºC (gray) for 24 additional hours (h) in the dark. For photobleaching quantification, seedlings were subsequently exposed to white light (WL) for 3 days. Data represent means ± SEM of biological replicates (t-test). a.u., arbitrary units. **(C)** Boxplot representations of the cotyledon aperture (left) and hypocotyl length (right) of at least 35 WT, *pif4* and *PIF4-L* seedlings grown as in (*B*). Mann Whitney test was used to define statistically differences. **(D)** Venn diagram showing overlap among heat-regulated genes in WT seedlings defined in this study and PIF-regulated and PIF-bound genes defined previously (Pfeiffer et al., 2014) (two-sided Fisher’s exact test). **(E)** Heat responsiveness (fold change; FC) in WT, *pif4* and *PIF4-L* for heat-regulated genes in *pif4* seedlings (*n*=3). Different letters denote statistically significant differences between genotypes by Dunn’s test (*P*<0.05). (A-C) Asterisks indicate statistically significant differences from *pif4* (\**P*<0.05, \*\**P*<0.01 and \*\*\**P*<0.001; n.s., not significant), and *n* the number of biological replicates.

To confirm the role of *PIF4* alternative splicing in regulating heat-induced responses, we generated transgenic plants expressing the short isoform of *PIF4* under the control of its endogenous promoter (*PIF4p::PIF4-S*) and evaluated their morphology in the dark under control temperature conditions. These transgenic lines (*PIF4-S*) showed higher *PIF4* expression levels than the corresponding WT control (Figure 4A), and in all three lines, the short isoform was the predominantly expressed variant (Figure 4B). We then conducted a phenotypic analysis of seedlings grown in the dark for 3 and 4 days at 21ºC. Interestingly, our results showed that enhanced production of the short isoform consistently promoted cotyledon opening, while changes in the hypocotyl length were not always detectable (Figure 4C). Thus, the cotyledon phenotype of these plants resembles that of WT plants exposed to heat stress (Figure 2C and 2D), linking the production of this isoform with heat-induced morphological adaptations. Notably, cotyledon opening in these transgenic plants at 21ºC is less pronounced than in heat-stressed plants (37ºC), indicating that the production of this isoform is not the unique mechanism underlying heat-induced cotyledon opening.

**Figure 4.**
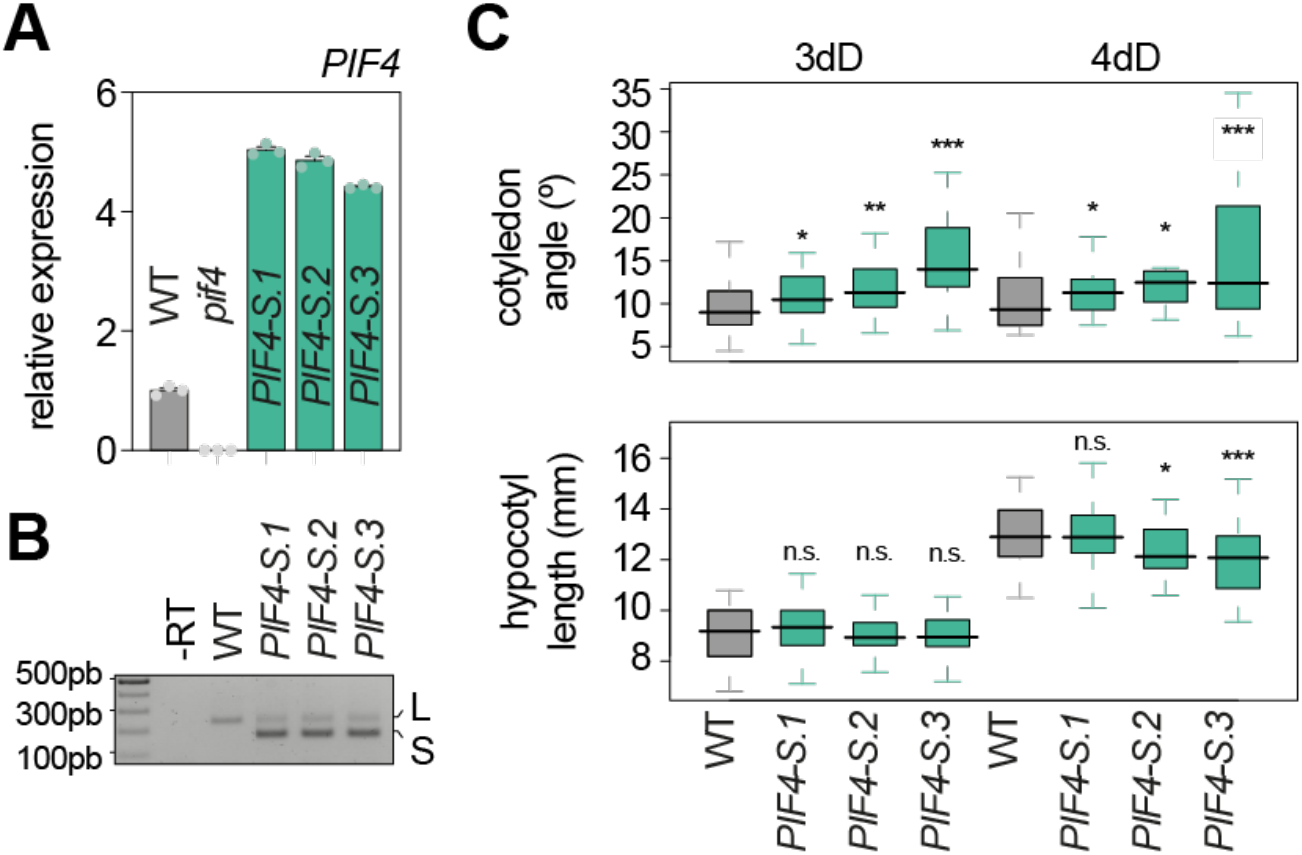
Enhancing expression of the *PIF4* short isoform promotes cotyledon opening in the dark. **(A)** RT-qPCR analysis of *PIF4* transcript levels in wild-type (WT), *pif4* and *PIF4-S* seedlings grown for 4 days in darkness. Values were normalized to *PP2A* and expression levels are expressed relative to the WT. Data represent means ± SEM of technical triplicates. **(B)** RT-PCR of the alternatively-spliced exon of *PIF4* in seedlings grown as in (*A*). -RT, no reverse transcriptase. **(C)** Boxplot representations of the cotyledon aperture (top) and hypocotyl length (bottom) of at least 28 WT, *pif4* and *PIF4-S* seedlings grown for 3 or 4 days (d) in the dark (D). Asterisks indicate statistically differences from WT at each day (Mann Whitney test; \**P*<0.05, \*\**P*<0.01 and \*\*\**P*<0.001; n.s., not significant).

## Discussion

Our study reveals, for the first time, that cotyledon opening is a developmental response of etiolated seedlings exposed to heat stress. Heat stress also exerts a repressive effect on hypocotyl elongation in etiolated seedlings, a phenomenon previously reported but not extensively studied (Hong and Vierling, 2000; Karayekov et al., 2013; Larkindale et al., 2005; Martín and Duque, 2022). Karayekov et al. linked this inhibitory effect on hypocotyls to altered functioning of light signaling components such as CONSTITUTIVE PHOTOMORPHOGENIC 1 (COP1), ELONGATED HYPOCOTYL5 (HY5) and circadian clock components. Notably, they also proposed a physiological rationale for this response: dark-germinated seedlings approaching the soil surface may become more susceptible to heat shock episodes due to their proximity to sunlight. They hypothesized that the molecular mechanisms triggered by these episodes would prime seedlings for imminent light exposure. Given that the first exposure to sunlight represents a critical phase for etiolated seedlings, requiring rapid adaptation to ensure survival, we agree that the early activation of photomorphogenesis-associated traits could be advantageous. Our finding that heat promotes cotyledon opening, another hallmark of photomorphogenesis (Arsovski et al., 2012), supports the hypothesis that, in etiolated seedlings, heat may function as a signal to induce photomorphogenesis. In addition, we show that heat enhances Pchlide accumulation in dark-grown seedlings. Although the conversion of Pchlide to chlorophyllide is light-dependent, Arabidopsis seedlings accumulate Pchlide in the dark to expedite this process upon light exposure (Sperling et al., 1997). However, the amount of Pchlide must be tightly balanced with the availability of the enzyme that catalyzes its conversion to prevent the generation of reactive oxygen species and subsequent cellular damage upon illumination (Mochizuki et al., 2010; Reinbothe et al., 1996). This raises the question of whether heat-induced Pchlide accumulation in etiolated seedlings is an adaptative mechanism to accelerate chlorophyll production and optimize the transition to autotrophic development, or whether it is a side effect of prematurely activating light signaling in the dark, an outcome that, as suggested by increased photobleaching, would negatively impact seedling survival.

Our data also indicate that the induction of these photomorphogenic traits in etiolated seedlings depends, at least in part, on a heat-specific regulatory event: the alternative splicing of *PIF4*. This represents a novel finding, as PIF4, despite being one of the most studied proteins in Arabidopsis, has been reported to be regulated only at the transcriptional and post-translational levels (Balcerowicz, 2020; Favero, 2020). Notably, our phenotypic analyses of seedlings with altered patterns of *PIF4* alternative splicing (*PIF4-L* and *PIF4-S*) suggest that the heat-induced isoform plays a role specifically in controlling cotyledon-related phenotypes. This implies that the mechanisms reported by Karayekov to control hypocotyl elongation under heat stress may operate in parallel with the alternative isoform of *PIF4*. Because our transcriptomic experiment was conducted using whole seedlings, we were unable to assess organ-specific effects in detail. We hypothesize that this alternative splicing event may be organ-specific or, alternatively, that the protein encoded by the heat-induced *PIF4* isoform may be preferentially active in cotyledons due to specific protein interactors and/or molecular targets. Further research into organ-specific dynamics is needed to elucidate why this alternative splicing event appears to predominantly affect cotyledon development. These findings would provide valuable insights into the organ-specific roles of PIF proteins, an emerging area of research (Dong et al., 2019; Sun et al., 2016; Zhang et al., 2021).

Our study focuses on the role of PIF4 under heat stress, a condition in which its function remains poorly understood. In fact, the few previous reports linking PIF function to heat stress have yielded contrasting results, with PIFs acting as either positive (Li et al., 2021) or negative (Yang et al., 2022) regulators of the response. Here, we significantly expand the current understanding of this transcription factor by demonstrating its involvement in heat stress and and further reinforcing its role as a key integrator of diverse environmental signals in the regulation of plant development (Paik et al., 2017). Importantly, modulating alternative splicing to alter isoform abundance is emerging as a promising strategy for developing stress-resilient plants (Alhabsi et al., 2025). In this context, investigating the role of the *PIF4* alternative isoform in heat-stressed adult plants, along with molecular strategies that specifically target the splicing sites involved in its regulation, could reveal a novel molecular target and offer an alternative genetic approach to enhance plant stress tolerance.

## Methods

### Plant materials

The *Arabidopsis thaliana pif1-1pif3-3pif4-2pif5-3* (*pifq*) and *pif4-2* mutant was obtained from the Nottingham Arabidopsis Stock Centre (NASC). *PIF4-L* transgenic plants expressing the *PIF4* coding region driven by the endogenous promoter, together with its respective *pif4* mutant control (*pif4-101*; Lorrain et al., 2008) were kindly provided by U. Pedmale (Cold Spring Harbor Laboratory, USA). *PIF4-S* transgenic plants were generated by PCR amplification and cloning a 1227-bp fragment containing the coding sequence region of the short splice variant under the control of a 2505-bp fragment upstream the PIF4 start codon corresponding to the PIF4 promoter, in the eGFP-tagged version of the binary pBA002 vector using the XbaI/AatI restriction sites (5’-GACGTT<u>TCTAGA</u>ATGGAACACCAAGGTTGGAG-3’and 5’-GT<u>GACGTC</u>CGAGTGGTCCAAACGAGAAC-3’). The pPIF4 promoter was insertved into the promoterless pBA002 via HindIII/XbaI restriction sites (5’-TGTGAAGCTTCCAAAGTAATAAAAGTTGCCACAAC-3’and 5’-GACGTTTCTAGAGTCAGATCTCTGGAGACATTTC-3’). The resulting constructs were introduced into *Agrobacterium tumefaciens* strain EHA105 and subsequently used for agroinfiltration-mediated transformation of Col-0 seedlings (Clough and Bent, 1998).

### Phenotypical and photobleaching analysis

Sterile seeds were sown on MS medium containing 1X Murashige and Skoog (MS) salts (Duchefa Biochemie), 2.5 mM MES (pH 5.7), 0.5 mM myo-inositol, and 0.8% agar (w/v). After stratification for 4 days at 4 ºC in darkness, seeds were subjected to a 3-hour light pulse to induce germination and then transferred to continuous darkness for 69 hours at 22 ºC. Maintaining the absence of light, seedlings were then either kept at 22 ºC for control conditions or transferred to 37 ºC for heat stress. Hypocotyl and cotyledon measurements of at least 30 seedlings and two biological replicates were carried out using the National Institutes of Health ImageJ software as described before (Sentandreu et al., 2011). Pictures were taken before and after exposure to stress as indicated in each figure. Photobleaching experiments were adapted from previous studies (Leivar et al., 2009), with 3-day-old etiolated seedlings being grown under control conditions or at 37 ºC for 24 hours and then transferred to continuous white light for 3 additional days (100 μmol·m^−2^·s^−1^). At this point, the percentage of seedlings that failed to become green were scored in three biological replicates.

### Protochlorophyllide quantification

Approximately 30 sterile seeds, sown on MS medium and stratified for 4 days at 4 ºC in the dark, were subjected to a 3-hour light pulse before being transferred to continuous darkness for 69 hours at 22 ºC. Seedlings were then either kept under these conditions (control) or transferred to 37 ºC in the dark to induce heat stress. Whole seedlings were collected in the dark 24 hours later, flash-frozen in liquid N_2_,and ground before extraction with 0.75 mL ice-cold 9:1 acetone:0.1 M NH4OH, as described previously (Terry and Kacprzak, 2019). The resulting mixture was vortexed for 1 minute and then centrifuged at 14,000 rpm at 4 ºC for 5 minutes. After supernatant recovery, the protochlorophyllide (Pchlide) content was determined as the peak value (635 nm) of the fluorescence emission spectrum between 600-700 nm, measured with a bandwidth of 5 nm after excitation at 440 nm and using a Synergy Neo2 microplate reader (Biotek). Pchlide data is shown as the average of Pchlide per seedling of four biological replicates.

### Gene expression and PSI quantification from RNA extraction

Total RNA was extracted from *Arabidopsis thaliana* seedlings using the InnuPREP Plant RNA kit (Analytik Jena BioSolutions) and 1 µg treated with DNase I to remove genomic DNA. cDNA synthesis using the oligo dT primer and the enzyme SuperScript III reverse transcriptase (Invitrogen) was conducted in the presence of RNase Out (Invitrogen). The cDNA was then used to quantify either gene expression or exon skipping of *PIF4*’s fifth exon. In both cases, three biological replicates were analyzed for each condition and/or genotype tested. Gene expression was measured by Reverse Transcription-quantitative PCR (RT-qPCR) using a QuantStudioTM 7 Flex Real-Time PCR System 384-well format and the Absolute SYBR Green ROX Mix (Thermo Scientific) on 2.5 µL of cDNA (diluted 1:10) per 10 µL of reaction volume, containing 300 nM of each gene-specific primer (see below). The *PP2A* gene was used for normalization (Shin et al., 2007). Exon skipping of the fifth exon of *PIF4* was quantified from RT-PCRs using primers spanning the two adjacent exons. These primer sequences were obtained from PastDB (Plant alternative splicing and transcription Data Base; www.pastdb.crg.eu; Martín et al., 2021). The resulting bands were quantified using the National Institutes of Health ImageJ software.

### RNA sequencing

RNA was extracted from 3-day-old WT, *pif4-101, PIF4-L*.*1* and *PIF4-L*.*2* etiolated seedlings grown for 3 hours at 37 ºC or 22 ºC in complete darkness. Oligo dT, non-strand specific libraries from triplicate biological replicates were built and sequenced using NextSeq500 at the Gulbenkian Institute for Molecular Medicine (GIMM). An average of 15 million 75-nucleotide single-end reads were generated per sample. Raw sequencing data was submitted to the Sequence Read Archive (accession number GSE200247).

### Gene expression quantification from RNA sequencing data

Quantification of *Arabidopsis thaliana* transcript expression from our RNA-seq experiment (GSE200247) and public sequencing data (Dataset S1) was performed using vast-tools v2.5.1 and v2.2.2 (Martín et al., 2021; Tapial et al., 2017), respectively. For each Arabidopsis transcript, this tool provides the number of mapped reads per million mapped reads divided by the number of uniquely mappable positions of the transcript (cRPKM; corrected-for-mappability reads per kbp of mappable sequence per million mapped reads) (Labbé et al., 2012). To identify genes differentially expressed between different temperatures, we used vast-tools compare_expr using the option -norm to perform a quantile normalization of cRPKM values between samples. Next, we filtered out the genes that were not expressed at cRPKM > 5 and had read counts > 50 across all the replicates of at least one of the samples compared. Finally, differentially-expressed genes were defined as those with a fold change of at least 2 between each of the individual replicates from each genotype. See https://github.com/vastgroup/vast-tools for details.

### PSI quantification from RNA sequencing data

We employed vast-tools v2.2.2 to quantify alternative splicing from public sequencing data (Martín et al., 2021; Tapial et al., 2017). This tool quantifies exon skipping (ES), intron retention (IR) and alternative donor (ALTD) and acceptor (ALTA) site choices. For all these types of events, vast-tools estimates the Percent Spliced In (PSI) of the alternative sequence using only exon-exon (or exon-intron for IR) junction reads and provides information about the read coverage See https://github.com/vastgroup/vast-tools for details. Data shown in Figure 1 and Supplemental Table 1 indicate the PSI quantification of specific alternative splicing events in the subfamily XV of the bHLH transcription factors (see below) using a wide array of samples (Supplemental Table 1).

## Data availability

RNA-seq data have been deposited in Gene Expression Omnibus (GEO) (GSE200247).

## Acknowledgments

We thank U. Pedmale for kindly providing *pif4-101* mutants and *PIF4pro:PIF4-3xFlag* transgenic lines, and V. Nunes for excellent plant care at the Gulbenkian Institute for Molecular Medicine (GIMM) Plant Facility. This work was funded by Fundação para a Ciência e a Tecnologia (FCT) through grants PTDC/BIA-FBT/31018/2017, PTDC/BIA-BID/30608/2017 and PTDC/ASP-PLA/2550/2021 as well as by the Spanish Ministry of Science and Innovation trough grant PID2021-125223NA-I00 (MCIN/AEI/10.13039/501100011033/FEDER). Funding from the research unit GREEN-it “Bioresources for Sustainability” (ID/04551/2025, UID/PRR/04551/2025) and the Generalitat de Catalunya (AGAUR, GRE2021, ref. SGR00873) is also acknowledged. G.M. was supported by an EMBO Long-Term Fellowship (ALTF 1576-2016), a Marie Skłodowska-Curie Individual Postdoctoral Fellowship (EU project 750469) and a Ramón y Cajal fellow from the Spanish Ministry of Science and Innovation (RYC2021-032539-I). T.L was supported by a Marie Skłodowska-Curie Individual Postdoctoral Fellowship (EU project 706274).

## Author contributions

M.N.-G., B.A., D.S., T.L. and G.M performed the experiments and analyzed the data. All authors discussed the results. G.M. conceived the project and designed research. G.M. and P.D. wrote the manuscript.

## Supplemental Information

**Supplemental File**. Supplemental Figures 1-10 and Supplemental Tables 1-2.

**Supplemental Table 1. List of publicly available RNA sequencing samples used**. Samples available at the Short Read Archive (SRA) were used to quantify the PSI (Percent Spliced In) and expression values shown in Figure 1, Supplemental Figure 2 and Supplemental Figure 3. Columns indicate our sample classification based on their growth conditions and tissue source, which are summarized in the sample description column, as well as the original SRA code number and name. For simplification PSI quantification was performed in samples originated from merging the biological replicates.

## Notes

### Competing Interest Statement

The authors have declared no competing interest.

## References

Alhabsi A, Ling Y, Crespi M, Reddy ASN, Mahfouz M. 2025. Alternative Splicing Dynamics in Plant Adaptive Responses to Stress. Annual Review of Plant Biology 76:687–717. DOI: 10.1146/annurev-arplant-083123-090055

Arsovski AA, Galstyan A, Guseman JM, Nemhauser JL. 2012. Photomorphogenesis. The Arabidopsis Book 10:e0147–e0147. DOI: 10.1199/tab.0147

Balcerowicz M. 2020. PHYTOCHROME-INTERACTING FACTORS at the interface of light and temperature signalling. Physiologia Plantarum 169:347–356. DOI: 10.1111/ppl.13092

Casson SA, Franklin KA, Gray JE, Grierson CS, Whitelam GC, Hetherington AM. 2009. phytochrome B and PIF4 Regulate Stomatal Development in Response to Light Quantity. Current Biology 19:229–234. DOI: 10.1016/j.cub.2008.12.046

Choi H, Oh E. 2016. PIF4 Integrates Multiple Environmental and Hormonal Signals for Plant Growth Regulation in <italic>Arabidopsis</italic>. Molecules and Cells 39:587–593. DOI: 10.14348/molcells.2016.0126

Clough SJ, Bent AF. 1998. Floral dip: a simplified method for Agrobacterium-mediated transformation of Arabidopsis thaliana. The Plant Journal 16:735–743. DOI: 10.1046/j.1365-313x.1998.00343.x

Dong J, Sun N, Yang J, Deng Z, Lan J, Qin G, He H, Deng XW, Irish VF, Chen H, Wei N. 2019. The Transcription Factors TCP4 and PIF3 Antagonistically Regulate Organ-Specific Light Induction of SAUR Genes to Modulate Cotyledon Opening during De-Etiolation in Arabidopsis. The Plant cell 31:1155–1170. DOI: 10.1105/tpc.18.00803

Favero DS. 2020. Mechanisms regulating PIF transcription factor activity at the protein level. Physiologia Plantarum 169:325–335. DOI: 10.1111/ppl.13075

Gangappa SN, Berriri S, Kumar SV. 2017. PIF4 Coordinates Thermosensory Growth and Immunity in Arabidopsis. Current Biology 27:243–249. DOI: 10.1016/j.cub.2016.11.012

Hong S-W, Vierling E. 2000. Mutants of Arabidopsis thaliana defective in the acquisition of tolerance to high temperature stress. Proceedings of the National Academy of Sciences 97:4392–4397. DOI: 10.1073/pnas.97.8.4392

Kan Y, Mu X-R, Gao J, Lin H-X, Lin Y. 2023. The molecular basis of heat stress responses in plants. Molecular Plant 16:1612–1634. DOI: 10.1016/j.molp.2023.09.013

Karayekov E, Sellaro R, Legris M, Yanovsky MJ, Casal JJ. 2013. Heat Shock– Induced Fluctuations in Clock and Light Signaling Enhance Phytochrome B– Mediated Arabidopsis Deetiolation. The Plant Cell 25:2892–2906. DOI: 10.1105/tpc.113.114306

Kim Sara, Hwang G, Kim Soohwan, Thi TN, Kim H, Jeong J, Kim Jaewook, Kim Jungmook, Choi G, Oh E. 2020. The epidermis coordinates thermoresponsive growth through the phyB-PIF4-auxin pathway. Nature Communications 11:1053. DOI: 10.1038/s41467-020-14905-w

Koini MA, Alvey L, Allen T, Tilley CA, Harberd NP, Whitelam GC, Franklin KA. 2009. High Temperature-Mediated Adaptations in Plant Architecture Require the bHLH Transcription Factor PIF4. Current Biology 19:408–413. DOI: 10.1016/j.cub.2009.01.046

Labbé RM, Irimia M, Currie KW, Lin A, Zhu SJ, Brown DDR, Ross EJ, Voisin V, Bader GD, Blencowe BJ, Pearson BJ. 2012. A Comparative Transcriptomic Analysis Reveals Conserved Features of Stem Cell Pluripotency in Planarians and Mammals. STEM CELLS 30:1734–1745. DOI: 10.1002/stem.1144

Larkindale J, Hall JD, Knight MR, Vierling E. 2005. Heat Stress Phenotypes of Arabidopsis Mutants Implicate Multiple Signaling Pathways in the Acquisition of Thermotolerance. Plant Physiology 138:882–897. DOI: 10.1104/pp.105.062257

Lee C-M, Thomashow MF. 2012. Photoperiodic regulation of the C-repeat binding factor (CBF) cold acclimation pathway and freezing tolerance in <em>Arabidopsis thaliana</em>. Proceedings of the National Academy of Sciences 109:15054– 15059. DOI: 10.1073/pnas.1211295109

Leivar P, Monte E. 2014. PIFs: Systems Integrators in Plant Development. The Plant Cell 26:56–78. DOI: 10.1105/tpc.113.120857

Leivar P, Monte E, Oka Y, Liu T, Carle C, Castillon A, Huq E, Quail PH. 2008. Multiple Phytochrome-Interacting bHLH Transcription Factors Repress Premature Seedling Photomorphogenesis in Darkness. Current Biology 18:1815–1823. DOI: 10.1016/j.cub.2008.10.058

Leivar P, Quail PH. 2011. PIFs: pivotal components in a cellular signaling hub. Trends in Plant Science 16:19–28. DOI: 10.1016/j.tplants.2010.08.003

Leivar P, Tepperman JM, Monte E, Calderon RH, Liu TL, Quail PH. 2009. Definition of Early Transcriptional Circuitry Involved in Light-Induced Reversal of PIF-Imposed Repression of Photomorphogenesis in Young <em>Arabidopsis</em> Seedlings. The Plant Cell 21:3535–3553. DOI: 10.1105/tpc.109.070672

Li B, Gao K, Ren H, Tang W. 2018. Molecular mechanisms governing plant responses to high temperatures. Journal of Integrative Plant Biology 60:757–779. DOI: 10.1111/jipb.12701

Li N, Bo C, Zhang Y, Wang L. 2021. PHYTOCHROME INTERACTING FACTORS PIF4 and PIF5 promote heat stress induced leaf senescence in Arabidopsis. Journal of Experimental Botany 72:4577–4589. DOI: 10.1093/jxb/erab158

Liao H, Feng B, Wen M, Du C, Zhong X, Lu Q, Gong G, Mo J, Huang H, Zhang S, Huang R. 2025. The transcription factors PIF4 and PIF5 interact with WRINKLED1 to modulate fatty acid biosynthesis during seed maturation. Plant Physiology 197:kiaf141. DOI: 10.1093/plphys/kiaf141

Liu Z, Zhang Y, Wang J, Li P, Zhao C, Chen Y, Bi Y. 2015. Phytochrome-interacting factors PIF4 and PIF5 negatively regulate anthocyanin biosynthesis under red light in Arabidopsis seedlings. Plant Science 238:64–72. DOI: 10.1016/j.plantsci.2015.06.001

Lorrain S, Allen T, Duek PD, Whitelam GC, Fankhauser C. 2008. Phytochrome-mediated inhibition of shade avoidance involves degradation of growth-promoting bHLH transcription factors. The Plant Journal 53:312–323. DOI: 10.1111/j.1365-313X.2007.03341.x

Lucyshyn D, Wigge PA. 2009. Plant Development: PIF4 Integrates Diverse Environmental Signals. Current Biology 19:R265–R266. DOI: 10.1016/j.cub.2009.01.051

Martín G, Duque P. 2022. Etiolated Hypocotyls: A New System to Study the Impact of Abiotic StressAbiotic stresses on Cell Expansion - Environmental Responses in Plants: Methods and Protocols. In: Duque P, Szakonyi D (Eds). Springer US. p. 195–205. DOI: 10.1007/978-1-0716-2297-1_13

Martín G, Márquez Y, Mantica F, Duque P, Irimia M. 2021. Alternative splicing landscapes in Arabidopsis thaliana across tissues and stress conditions highlight major functional differences with animals. Genome biology 22:35. DOI: 10.1186/s13059-020-02258-y

Mochizuki N, Tanaka R, Grimm B, Masuda T, Moulin M, Smith AG, Tanaka A, Terry MJ. 2010. The cell biology of tetrapyrroles: a life and death struggle. Trends in Plant Science 15:488–498. DOI: 10.1016/j.tplants.2010.05.012

Nicolas M, Rodríguez-Buey ML, Franco-Zorrilla JM, Cubas P. 2015. A Recently Evolved Alternative Splice Site in the <em>BRANCHED1a</em> Gene Controls Potato Plant Architecture. Current Biology 25:1799–1809. DOI: 10.1016/j.cub.2015.05.053

Paik I, Kathare PK, Kim J-I, Huq E. 2017. Expanding Roles of PIFs in Signal Integration from Multiple Processes. Molecular plant 10:1035–1046. DOI: 10.1016/j.molp.2017.07.002

Penfield S, Josse E-M, Halliday KJ. 2010. A role for an alternative splice variant of PIF6 in the control of Arabidopsis primary seed dormancy. Plant Molecular Biology 73:89–95. DOI: 10.1007/s11103-009-9571-1

Pfeiffer A, Shi H, Tepperman JM, Zhang Y, Quail PH. 2014. Combinatorial Complexity in a Transcriptionally Centered Signaling Hub in Arabidopsis. Molecular Plant 7:1598–1618. DOI: 10.1093/mp/ssu087

Pham VN, Kathare PK, Huq E. 2018. Phytochromes and Phytochrome Interacting Factors. Plant Physiology 176:1025–1038. DOI: 10.1104/pp.17.01384

Quint M, Delker C, Franklin KA, Wigge PA, Halliday KJ, van Zanten M. 2016. Molecular and genetic control of plant thermomorphogenesis. Nature Plants 2:15190. DOI: 10.1038/nplants.2015.190

Reinbothe S, Reinbothe C, Apel K, Lebedev N. 1996. Evolution of Chlorophyll Biosynthesis – The Challenge to Survive Photooxidation. Cell 86:703–705. DOI: 10.1016/S0092-8674(00)80144-0

Sakuraba Y, Jeong J, Kang M-Y, Kim J, Paek N-C, Choi G. 2014. Phytochrome-interacting transcription factors PIF4 and PIF5 induce leaf senescence in Arabidopsis. Nature Communications 5:4636. DOI: 10.1038/ncomms5636

Sentandreu M, Martín G, González-Schain N, Leivar P, Soy J, Tepperman JM, Quail PH, Monte E. 2011. Functional profiling identifies genes involved in organ-specific branches of the PIF3 regulatory network in Arabidopsis. The Plant Cell 23:3974–3991. DOI: 10.1105/tpc.111.088161

Seo PJ, Kim MJ, Ryu J-Y, Jeong E-Y, Park C-M. 2011. Two splice variants of the IDD14 transcription factor competitively form nonfunctional heterodimers which may regulate starch metabolism. Nature Communications 2:303. DOI: 10.1038/ncomms1303

Shin J, Kim K, Kang H, Zulfugarov IS, Bae G, Lee C-H, Lee D, Choi G. 2009. Phytochromes promote seedling light responses by inhibiting four negatively-acting phytochrome-interacting factors. Proceedings of the National Academy of Sciences 106:7660–7665. DOI: 10.1073/pnas.0812219106

Shin J, Park E, Choi G. 2007. PIF3 regulates anthocyanin biosynthesis in an HY5-dependent manner with both factors directly binding anthocyanin biosynthetic gene promoters in Arabidopsis. The Plant Journal 49:981–994. DOI: 10.1111/j.1365-313X.2006.03021.x

Sperling U, van Cleve B, Frick G, Apel K, Armstrong GA. 1997. Overexpression of light-dependent PORA or PORB in plants depleted of endogenous POR by far-red light enhances seedling survival in white light and protects against photooxidative damage. The Plant Journal 12:649–658. DOI: 10.1046/j.1365-313X.1997.00649.x

Sun N, Wang J, Gao Z, Dong J, He H, Terzaghi W, Wei N, Deng XW, Chen H. 2016. Arabidopsis SAURs are critical for differential light regulation of the development of various organs. Proceedings of the National Academy of Sciences of the United States of America 113:6071–6076. DOI: 10.1073/pnas.1604782113

Tapial J, Ha KCH, Sterne-Weiler T, Gohr A, Braunschweig U, Hermoso-Pulido A, Quesnel-Vallières M, Permanyer J, Sodaei R, Marquez Y, Cozzuto L, Wang X, Gómez-Velázquez M, Rayon T, Manzanares M, Ponomarenko J, Blencowe BJ, Irimia M. 2017. An atlas of alternative splicing profiles and functional associations reveals new regulatory programs and genes that simultaneously express multiple major isoforms. Genome Research 27:1759–1768.

Terry MJ, Kacprzak SM. 2019. A Simple Method for Quantification of Protochlorophyllide in Etiolated Arabidopsis Seedlings BT - Phytochromes: Methods and Protocols. In: Hiltbrunner A (Ed). Springer New York. p. 169–177. DOI: 10.1007/978-1-4939-9612-4_14

Toledo-Ortiz G, Huq E, Quail PH. 2003. The Arabidopsis basic/helix-loop-helix transcription factor family. The Plant Cell 15:1749–1770. DOI: 10.1105/tpc.013839

Wang Q, Zhao L, Shao T, Xu Z, Zhu Z. 2025. Salt-responsive SSN1 condensation in nucleus facilitates PIF4 degradation to regulate Arabidopsis salt tolerance. The Plant Journal 123:e70389. DOI: 10.1111/tpj.70389

Yang J, Qu X, Ji L, Li G, Wang Chen, Wang Changyu, Zhang Y, Zheng L, Li W, Zheng X. 2022. PIF4 Promotes Expression of HSFA2 to Enhance Basal Thermotolerance in Arabidopsis. International Journal of Molecular Sciences. DOI: 10.3390/ijms23116017

Zhang Y, Li N, Wang L. 2021. Phytochrome interacting factor proteins regulate cytokinesis in Arabidopsis. Cell Reports 35:109095. DOI: 10.1016/j.celrep.2021.109095

